# ANUBI: A Platform for Affinity Optimization of Proteins and Peptides in Drug Design

**DOI:** 10.1101/2025.10.20.683353

**Authors:** Damiano Buratto, Wanding Wang, Xinyi Zhang, Qiujie Zhu, Jia Meng, Daniel J. Rigden, Ruhong Zhou, Francesco Zonta

## Abstract

The increasing availability of computational power is opening unprecedented opportunities in computational biology and drug design. Computer simulations based on physical models can now reproduce or replace critical biophysical experiments such as binding affinity evaluations in the drug screening process. Here we present ANUBI (ANUBI Nexus for Understanding Binding Interactions), a software package that automates sequence space exploration and binding free energy calculations to optimize protein or peptide drug candidates for improved target binding. Starting from a user-provided molecular model of the drug-target interaction, ANUBI systematically evaluates point mutations in selected regions using Monte Carlo methodology, retaining favorable mutations based on calculated binding affinity differences. We demonstrate that this approach efficiently samples sequence space, generating dozens of optimized variants in timeframes comparable to experimental approaches at substantially reduced cost. As proof of concept, we applied ANUBI to an antibody-antigen complex and a peptide-protein interaction, identifying variants with significantly improved predicted binding energy (approximately 20 kcal/mol, calculated using the MMPBSA method), within 20 days of computation on a single GPU.

TOC GRAPHIC

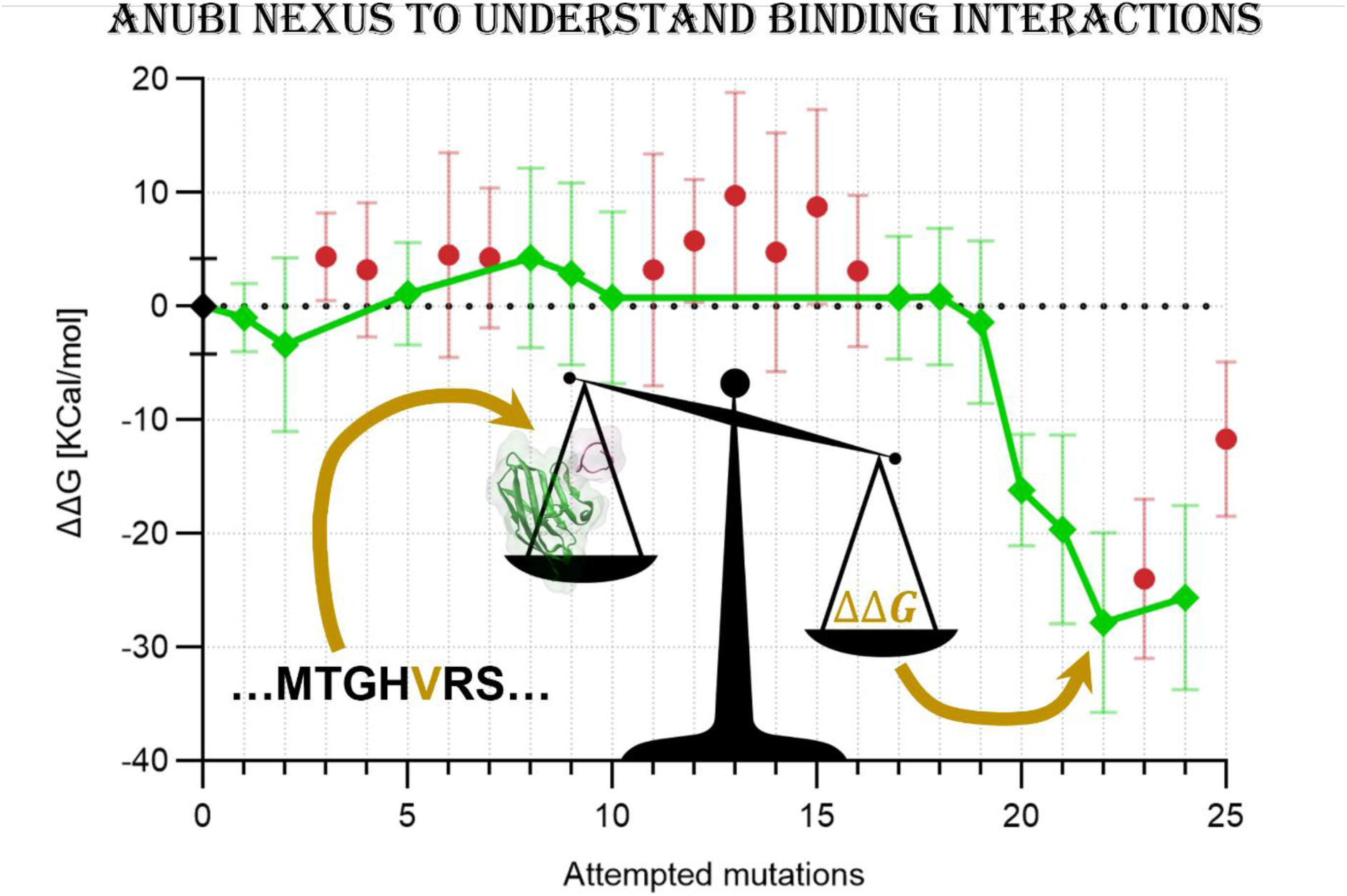

## INTRODUCTION

Despite recent progress in structure prediction and design, modeling protein-ligand or protein-protein interactions and evaluating their binding affinity still represent major challenges in theoretical biochemistry ^1,2^. Any method that allows rapid and accurate evaluation of binding affinity between molecules, particularly proteins, would represent a major milestone in drug design, as binding affinity evaluation is the first critical step in the drug screening process ^3^.

Simplified scoring functions, widely implemented in programs such as AutoDock Vina, Schrödinger Suite, and Rosetta ^4–6^, have achieved reasonable success in screening small molecule libraries against protein targets. Machine learning methods are also emerging, showing remarkable accuracy in the prediction of binding affinities of small molecules to their targets ^7^. These methods can rapidly evaluate thousands of compounds and identify promising lead candidates, making them valuable tools in early-stage drug discovery.

However, similar approaches are considerably less successful when applied to protein-protein or protein-peptide interactions. While machine learning methods ^8–11^ and traditional docking programs ^12–14^ can produce reliable structural predictions of protein complexes, accurately evaluating binding affinities remains challenging. An impressive breakthrough in this sense has been achieved recently thanks to generative AI methods such as RosettaFold and BindCraft ^15,16^ which are able to generate protein sequences displaying high binding affinity to a desired target.

Despite these advances, all current methodologies share a fundamental limitation since the scoring of possible binders is based on a single representation of the docked complex. Machine learning methods, for example, use confidence scores like pLDDT (predicted Local Distance Difference Test, the model’s confidence in predicting the position of each residue) as proxies for binding quality, despite these metrics reflecting structural plausibility rather than thermodynamic stability. Biological macromolecules, instead, contain flexible regions that can sample multiple configurations, and even minor side chain rearrangements can result in significant energetic differences that simplified scoring functions fail to capture adequately^17^. The issue is particularly important in dealing with macromolecules with low structural organizations, such as peptides, intrinsically disordered proteins, or short RNAs.

Molecular Dynamics (MD) simulations can overcome this limitation, at least in principle. While their computational cost is very high compared to simplified methods, they have become accessible to average users in the field due to methodological improvements and the exponential increase in computing power over time. In previous work ^18,19^, we demonstrated that MD-based methodologies can be used for screening biological drugs (such as antibodies) when paired with smart sampling of the vast chemical space.

Based on such experience, here we propose ANUBI (ANUBI Nexus for Understanding Binding Interactions), a revised and improved version of the pipeline introduced in ^18^, which aims to screen biological drugs (peptides or proteins) based on their binding affinity to a target. Like the Egyptian god Anubis, who weighed hearts to determine worthiness, ANUBI evaluates the binding “weight” of candidate molecules to determine which merits advancement in the drug discovery pipeline.

The major advantages of the ANUBI platform include (i) its automated workflow requiring minimal computational expertise, (ii) efficient Monte Carlo-based sequence sampling, (iii) adaptability to both protein and peptide optimization, and (iv) computational efficiency enabling evaluation of hundreds of sequences within weeks, making ANUBI a viable alternative to purely experimental workflows. While ANUBI is designed for optimization rather than de novo design, it can refine initial configurations obtained from experimental structures or from generative AI methodologies such as RosettaFold, ProteinMPNN, or BindCraft, thereby improving their success rate and reducing experimental screening burden.

Compared to our previous work ^18^, ANUBI introduces significant methodological advances: (i) we have consolidated the sequence sampling method by systematically validating the efficacy of then Monte Carlo algorithm across different targets with hundreds of independent runs, (ii) we have created an accessible and documented Python package for community use, and (iii) we have implemented and validated a peptide optimization mode, expanding applicability to the rapidly growing field of therapeutic peptides ^20^.

In the following sections, we describe the implementation details, validate the approach on two representative systems, and discuss the platform’s potential applications in drug design.

## IMPLEMENTATION

### ANUBI Workflow

ANUBI is a user-friendly and fully automated computational framework developed for the in-silico optimization of the binding affinity of protein drugs (e.g., antibodies) or peptides to a desired target.

It is composed of two main modules: the *Sequence sampling module* generates the amino acid sequence of the drug candidate, while the *Energy module* evaluates the binding energy of the candidate to the target. Through an automatic workflow (**Figure 1**), ANUBI combines the two modules to iteratively generate new candidates and produce sequences with improved theoretical binding affinities.

**Figure 1.**
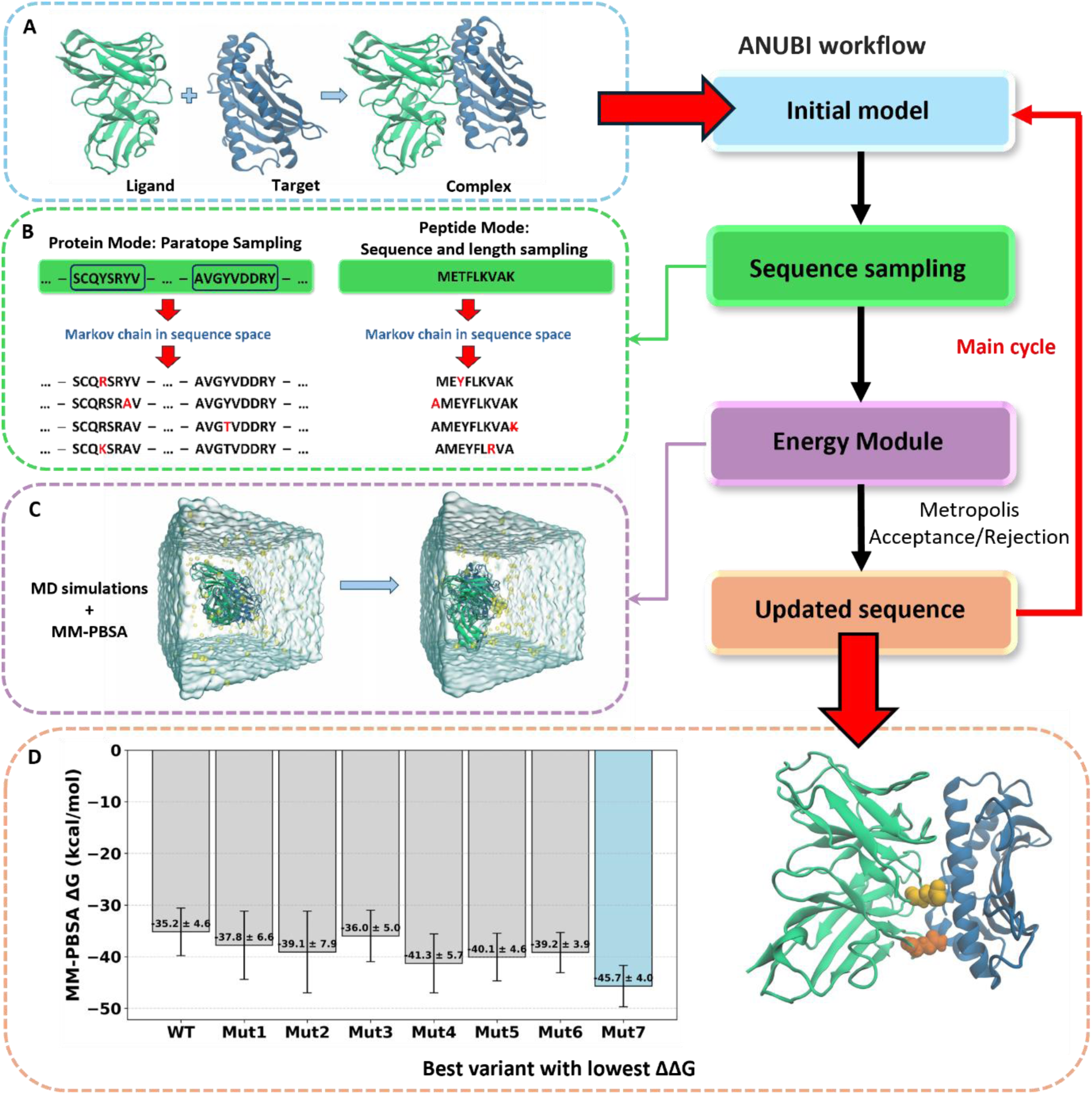
Overview of ANUBI. **(A)** The pipeline begins with the wild-type protein–ligand complex provided in PDB format. **(B)** Following these steps, modifications to the sequence of the ligand are proposed by the Monte Carlo algorithm. This includes amino acid substitution in the paratope region, and, in peptide mode, addition of an Alanine or removal of a residue at the N-term or C-term. **(C)** The proposed Monte Carlo move is accepted or rejected according to the Metropolis algorithm based on binding energy calculations. **(D)** The cycle is iterated many times until a new sequence leading to the best binding affinity is given as output. Mutations leading to the best binders are highlighted as an example.

In its current implementation, the sequence sampling is done with a Monte Carlo Metropolis algorithm, aiming to generate sequences with better (i.e., more negative) binding free energy. The binding energy is evaluated using MD simulations paired with the MMPBSA method ^21^. Each module operates independently, allowing flexibility and scalability across different protein-ligand systems. We plan to implement alternative methodologies for both modules in the future, to allow more freedom of choice to the user.

### Input files

The software requires a user-supplied PDB file (‘*complex.pdb*’) with the initial model of the ligand-target complex. This model needs to be pre-relaxed and equilibrated through MD simulations. Users need to provide stable and accurate input structures, as models with structural errors, such as missing atoms and amino acids or presenting major steric clashes, may result in the incorrect execution of the following pipeline. The use of AMBER-style PDB format is recommended to ensure consistency in atom and residue naming, which facilitates correct force field assignment and avoids preprocessing errors.

Model preparation requires a certain level of expertise in molecular modelling and cannot be automated in a straightforward way. Nevertheless, it is still a relatively low entry barrier since model generation, energy minimization, and equilibrium MD simulations are widely used tools. Every other step in the follow-up pipeline does not require human intervention. Only the coordinates of the atoms belonging to the proteins are required, as the generation of the periodic boundary condition box, solvation, and addition of ions are done automatically at each iteration.

In addition to the initial model, ANUBI requires a user-defined YAML configuration file (*‘infile.yaml’*) to specify the required input files, computational environment, and key parameters for different modules.

Altogether, the input for the current implementation of ANUBI looks as it follows:

1. **Starting target-ligand configuration**. User-provided structure files (PDB format) representing the ligand-target complex.
2. **Basic Settings**. Paths to the required software components: Conda environment activation script, VMD, GROMACS, and Conda environment names for *gmx_mmpbsa* and Modeller (see below for a detailed explanation of each component).
3. **Free Energy Calculation settings**. Definition of the receptor and ligand, and other relevant settings, e.g., which part of the trajectory will be used for the MMPBSA calculations.
4. **Sequence sampling Settings**. List of residues that are allowed to be mutated by the Monte Carlo and the total length of the Monte Carlo sampling cycles.
5. **Run Configurations**. Specific parameters (such as CPU numbers) required for optimizing parallel performance.

This structured configuration file ensures method flexibility and results reproducibility, while simplifying the setup process across different design tasks. Users can easily customize the parameters to fit their specific research needs and computational environment.

### Sequence sampling module

The aim of the pipeline is to generate protein or peptide candidate binders to a desired target after an exhaustive sampling of the sequence space. With 20 possible choices of standard amino acids, the chemical space becomes extremely large even when we allow mutations only in a few selected positions. To bypass this issue, we need to apply effective strategies to sample the sequence space.

In the current implementation of ANUBI, we choose a standard Monte Carlo Method for this purpose. As we showed in our previous work, this methodology is surprisingly effective and can produce improved sequences in a small number of iterations ^18^.

As proteins and peptides are biochemically distinct entities - differing in size, structural complexity, and conformational stability ^22^ - we decided to differentiate the *Protein Mode* from the *Peptide Mode* when dealing with sequence sampling (**Figure 2**).

**Figure 2.**
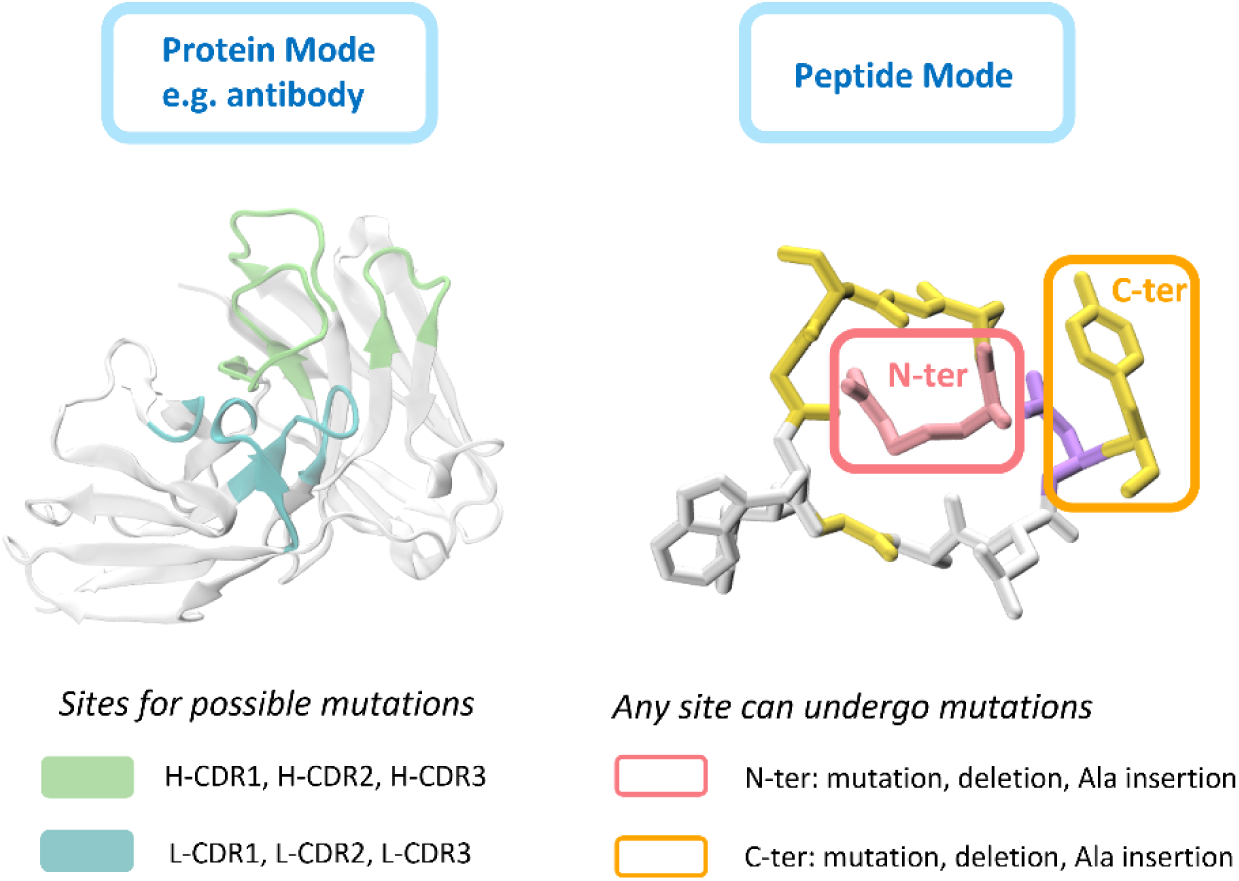
Sequence modification strategies in ANUBI. **(A)** Protein Mode introduces single-point mutations in the paratope region (or other part of the protein chosen by the user), for example the CDRs loops of an antibody, which are critical to the binding to the antigen. **(B)** Peptide Mode applies point mutations across the entire peptide sequence and additionally varies the sequence length by adding or removing residues at the N-terminal or C-terminal ends.

The Monte Carlo proposal move in the protein mode allows for a single point mutation of a residue within a user-selected list of residues (for example, the paratope region or the Complementary Determining Regions – CDRs – for an antibody). The list of residue positions for mutation is specified by the user in the configuration file (‘*infile.yaml*’). At each iteration, one residue is randomly selected from the defined list and mutated to a randomly chosen amino acid. The new amino acid is chosen from 17 standard residues, excluding cysteine, glycine, and proline. Cysteine is avoided because it can form non-native disulfide bonds that lead to a rearrangement of the local structure ^23^. Glycine, with its high backbone flexibility, tends to increase local conformational entropy and reduce stability. Proline’s rigid cyclic structure often causes kinks that disrupt helices and strands. Recent computational work also shows that modelling glycine and proline substitutions is challenging due to their effects on backbone geometry ^24^. By excluding these residues, the mutations are more likely to preserve structural stability and produce reliable results.

In peptide mode, in addition to the previously described single point mutations, we also allow for the change of the peptide length, which may happen through deletion of the first or last residue or insertion of an Alanine at the termini. The choice of Alanine as the insertion residue is due to its small, nonpolar methyl side chain, which introduces minimal steric hindrance while maintaining the flexibility of the peptide backbone ^25^.

The application of length-modifying operations is attempted after a user-defined number of attempted single points mutations, typically every 5 to 10 proposed moves, at which point the software randomly choses one of the four possibility (deletion at the N-terminus, deletion at the C-terminus, addition of an Ala at the N-terminus or addition of an Ala at the C-terminus).

Furthermore, to keep the peptide length under control, the user can define the minimum and maximum allowed lengths tailored to their projects. Generally, we expect to work with peptide lengths from 10 to 20 amino acids, as this is a typical range for bioactive and therapeutic peptides ^26,27^. Some studies, indeed, have reported that binding affinity tends to peak when peptides are approximately 18 to 20 amino acids in length, while further increases in length may lead to reduced affinity ^28^. In contrast, very short peptides are generally more soluble in water, but they often lack the structural support and interaction surface necessary for stable and specific binding ^22^.

The acceptance probability (*P*) of the proposed move (mutation, addition, or deletion of a residue) is determined according to the Metropolis algorithm: *P* = min {1, *e*^−*β*ΔΔ*G*^}, where ΔΔ*G* is the difference of the binding free energy before and after the mutation (see next section). In this way, mutations which improve the binding affinity (ΔΔ*G* < 0) are always accepted, but it is also possible to accept mutations with worse binding affinity, allowing the algorithm to escape local minima. The parameter *β* in the Metropolis acceptance criterion *e*^−*β*ΔΔ*G*^ functions analogously to inverse temperature scaled by the gas constant (1/*RT*) but is independent of the MD simulation temperature and can be tuned by the user to increase or decrease the probability of accepting configuration with worse binding affinity. In the different runs we used values around *β* = 2.0 *kcal*/*mol*. This was empirically determined to offer a good balance between exploration and convergence. In our tests, values *β* < 1 *kcal*/*mol* led to an overly conservative search in which almost no energetically unfavorable mutations are accepted, while values *β* ≥ 4 *kcal*/*mol* resulted in excessive acceptance of worse-scoring variants.

If the proposed move is accepted, ANUBI will update the current structural model by including the new mutation, and this will be the input for the next Monte Carlo cycle. Otherwise, if the mutation is not accepted, the structural model is not updated, and the next cycle will still use the unchanged structure as input.

This iterative process continues until the specified number of cycles is completed. As we will show later, this algorithm is able to find new sequences with improved binding affinity after 10 - 15 steps but tend to get stuck after 20 - 30 steps. In all our tests, different Monte Carlo runs rapidly diverged in sequence space. Therefore, to increase the sequence sampling, it is preferable to run numerous short instances of ANUBI, instead of increasing the length of a particular run.

### Automatic generation of molecular models after sequence modification

Binding free energies can be calculated in many ways, starting from a structural model of the interacting molecules. To obtain such a model after the changes proposed by the Sequence sampling module, we use *Modeller* software ^29^ starting from the previous structure as template.

This step typically imposes very minor changes to the structure and for this reason local strain and steric clashes introduced by the change can easily be eliminated through a two-step energy minimization protocol, which first involves the mutated residue itself and subsequently nearby residues in space.

In peptide mode, when an alanine residue is added to the one of the termini, local structural refinement is carried out to ensure smooth integration with the existing backbone. Conversely, when removing a terminal residue, the first or last residue is simply deleted without further adjustment, as such changes are generally well-tolerated and do not significantly affect structural integrity at the termini.

After the mutation step, the topology file for the new molecular system is created using the Gromacs suite (*pdb2gmx*), and a standard box for periodic boundary conditions is created. Water and ions (in a neutralizing solution of 0.15 M KCl), are added again at this stage as they are not retained during the mutation step. This fully solvated system then serves as input for binding free energy calculations, which determine whether the proposed mutation will be accepted or rejected.

### Energy module

As we have seen, acceptance or rejection of the proposed mutations is based on the binding free energies of the ligands to the target. More precisely, we define the binding free energy Δ*G* as:

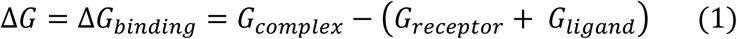

Were *G*_*receptor*_, *G*_*ligand*_ and *G*_*complex*_are, respectively, the Gibbs free energy of the receptor alone, the ligand alone, and the complex of the receptor and the ligand after the binding. As the receptor and the ligand should bind spontaneously, we expect Δ*G* to be negative. ΔΔ*G* represents the difference of Δ*G* after and before the proposed mutation:

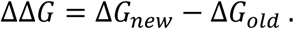

A negative value of ΔΔ*G* indicates that the new Δ*G* is more negative, i.e. the binding between the two molecules is stronger.

In the current implementation of ANUBI, binding free energies are calculated using the MMPBSA ^21^ method, and in particular the *gmx_mmpbsa* module ^30^. This is not the only possible choice, and we are planning to include additional methods in the future, in particular, FEP calculations ^31^. The reasons to prioritize MMPBSA methods for the initial release are due to two different advantages: (i) it provides absolute estimates of the binding energy, and (ii) it is relatively faster. However, it also comes with shortcomings because it does not evaluate the entropic contribution to the binding free energy Δ*G*, and because the estimates produced with this method have very high variance, even with small rearrangements of the atoms in the interface between the two molecules.

However, both these shortcomings can be attenuated by a careful sampling of the configuration space. This is done using MD simulations. We have shown that for computational efficiency it is preferable to run a series of short (<10 ns) MD trajectories and calculate the binding energy on hundreds of configurations extracted from these trajectories. In this way, all energy terms in equation (1) are computed as ensemble averages over selected MD frames ^19^.

These considerations lead to the following workflow (illustrated in **Figure 3**) for calculating the binding energies in ANUBI. The system is pre-processed as explained in the previous section and relaxed with a 1 ns equilibration in the NVT ensemble. Following this, ten independent 5 ns MD simulations are performed in the NPT ensemble. For each simulation, 150 different configurations are extracted from the last 3 ns of the trajectory and the binding energy for each of them is calculated using the MMPBSA method.

**Figure 3.**
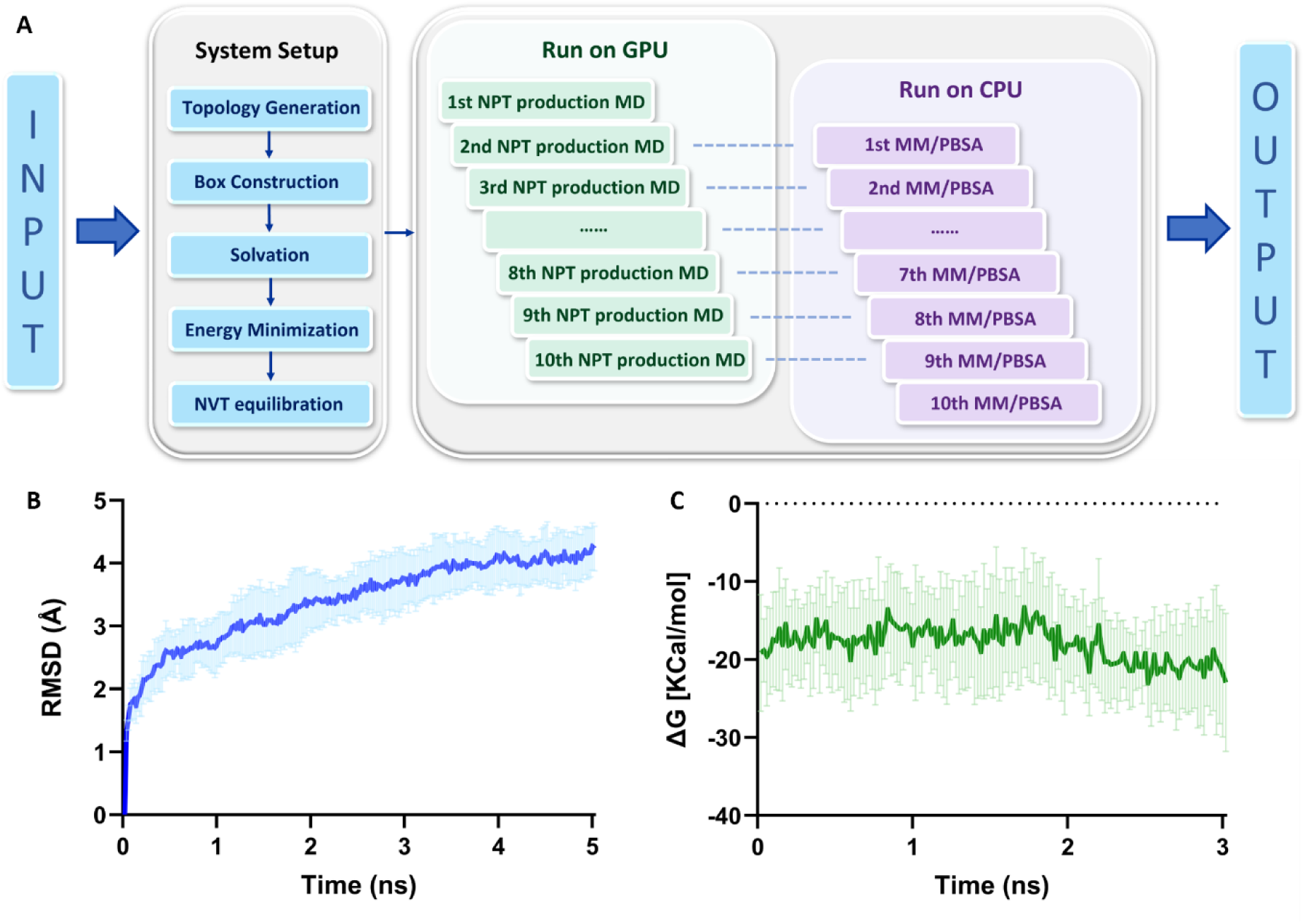
Overview of Energy Module for binding free energy calculation. **(A)** The module takes as input the original model or the modified model generated by Modeller and provide as output the binding free energy, which will be evaluated by the Monte Carlo algorithm. Setup steps (topology generation, box construction, water and ions addition, energy minimization,1 ns NVT equilibration) and energy calculation is done automatically. The binding energy is calculated using the MMPBSA algorithm along 10 different 5 ns NPT production runs MD simulations. MD simulations are run on the GPUs, while the energy calculation is run in parallel on the CPU. **(B)** Average and standard deviation of the RMSD and **(C)** average and standard deviation of binding energy calculated along the 10 different simulations. The binding energy is calculated as time average along the last 3 ns.

To improve computational efficiency, the MMPBSA calculations are executed in parallel with the subsequent NPT production simulations. Specifically, while the second production MD run is performed on the GPU, the MMPBSA analysis for the first production MD is simultaneously carried out on the CPU and analogously for the other replicas. This parallel design significantly reduces the total computation time and enables scalable execution of multiple replicates.

The average binding energy of the 1500 configurations obtained from the ten MD trajectories is then passed as result to the Monte Carlo cycle that will decide whether to accept and update the new configuration or reject it and go back to the previous configuration.

## RESULTS AND DISCUSSION

In this section, we provide two examples of applications of ANUBI in designing antagonist antibodies or peptides versus targets of interest. We want to demonstrate how ANUBI can be used as a computational tool to screen for candidates that will be passed on to the experimental pipeline. Therefore, we provide only evidence of the consistency and efficiency of the computational method. Experimental validation is ongoing for both targets, but it is out of the scope of the work presented here.

### Connexin 43 hemichannels

The first example is an antibody against a Cx43 hemichannel. We have previously designed and characterized AbEC1.1, an antibody against hemichannels composed of beta connexin sub-family members (Cx26, Cx30, and Cx32 ^32–34^). This antibody can also weakly bind and partially block Cx43 hemichannels. In our previous work, we attributed the difference in binding affinity to two amino acid differences between the beta connexin and Cx43 in the epitope region ^33^.

Here, we show how it is possible to adapt AbEC1.1 to Cx43 using ANUBI. The starting model is the previously published and characterized model of interaction between AbEC1.1 and the Cx26 hemichannels, where the two critical amino acids have been substituted (Q56L in the first extracellular loop and H176N in the second extracellular loop) with the corresponding Cx43 amino acids.

To improve the speed of simulation, only relevant parts are kept: the extracellular loops of the connexin hemichannel and a few other amino acids of the transmembrane regions proximal to such loop, and the variable domains of the antibody. To preserve the structural integrity of this truncated system, heavy atoms in the transmembrane regions were restrained using a harmonic potential (see Methods and **Figure 4 panel A**).

**Figure 4.**
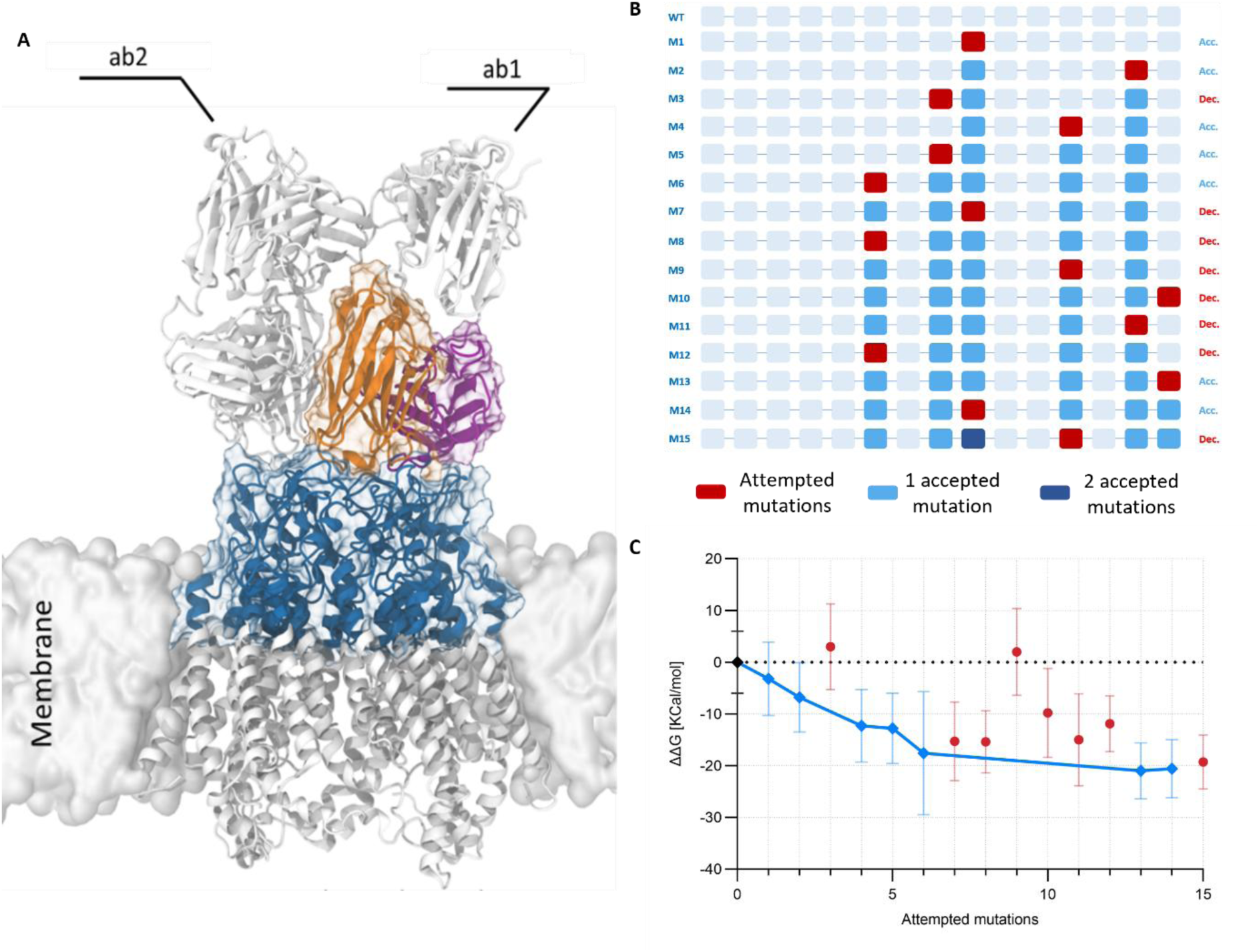
Affinity maturation in protein mode. The figure shows an example of a Monte Carlo run for the affinity maturation of an antibody against Cx43 hemichannels. **(A)** The model used for simulations is shown in color and was extracted from a previous model (in white) of two antibodies bound to the Cx43 hemichannel (see methods), to reduce the computational cost. **(B)** Schematic representations of the positions of attempted and accepted mutations. Each square represents an amino acid belonging to the paratope. The red squares represent sites where mutation is attempted, and darker blue squares represent accepted mutations. Acc. and Dec. indicate whether the mutation is accepted or declined at that specific step. **(C)** Graph of the binding energy difference along the simulation. Blue points represent the binding energy difference between the wild type and the mutations that have been accepted. The bars indicate the standard error of the computational results.

ANUBI is then used to adapt the antibody to the Cx43 hemichannels. **Figure 4 Panels B and C** show a single run of the optimization process. In this run, we allow for mutations only of key residues of the hCDRs 1 and 3 of the antibody. We can notice how the computed binding energy quickly decreases as new sequences are accepted by the Monte Carlo algorithm. We can also observe that mutations are accepted in several positions in the CDRs of the antibody, and at the end, six out of fifteen amino acids have been mutated, meaning that the final antibody can be considered completely different from AbEC1.1.

Most of the accepted mutations account for small *ΔΔG* values, but some produce large changes, providing valuable insight into the most critical positions for binding that can guide further optimization. In the example run of **Figure 4**, the total difference in *ΔΔG* for the best candidate of the Monte Carlo cycle, evaluated by the MMPBSA method, is predicted to be around 21 ± 5 kcal/mol. While this absolute value is unrealistic due to MMPBSA limitations, our previous work and ongoing experimental validations demonstrate that sequences showing significant computational improvements (>15 kcal/mol) translate into better binders also in experiments. The method’s value lies in rank-ordering candidates rather than providing quantitatively accurate binding energies.

### T cell immunoreceptor with Ig and ITIM (TIGIT)

To test the peptide mode, we chose as a target the T cell immunoreceptor with Ig and ITIM (TIGIT). Experimental validation and detailed methods of the design of the initial peptide will be given in a follow-up work. Here we can mention that different peptide binders were designed starting from known ligands of TIGIT (PVR: the poliovirus receptor protein, and MG1131: an antibody antagonist ^35,36^). The more promising candidate peptide was chosen to be carried on in the follow-up protocol, based on the RMSD of the whole system (around 4.0 Å) and a good initial binding affinity (-20 ± 6 kcal/mol, calculated with MMPBSA method), which assured that the peptide was stably bound to TIGIT during the simulations. Besides single point mutations, ANUBI attempted to modify the length of the peptide every 5 moves, by choosing with equal probability whether to add an alanine at one of the termini, or to delete one of the termini. The limits for the length of the peptide were set between the minimum of 8 and the maximum of 20 amino acids.

Like the antibody case presented above, we observe that several amino acid substitutions were accepted, and the final peptide maintained only four of the ten initial amino acids (**Figure 5**). Furthermore, the peptide changed its length from 10 to 12 amino acids, as two alanines were added during the affinity maturation process, one for each terminus, showing that length changes can be beneficial to adapt the peptide to the target. As previously observed, some substitutions are clearly more impactful, revealing hotspots for peptide design. The change of ΔΔ*G* evaluated with the MMPBSA from the beginning to the end of the cycle is 26 ± 8 Kcal/mol. As mentioned before, such a large value is unrealistic, and we expect to see less difference in experiments.

**Figure 5.**
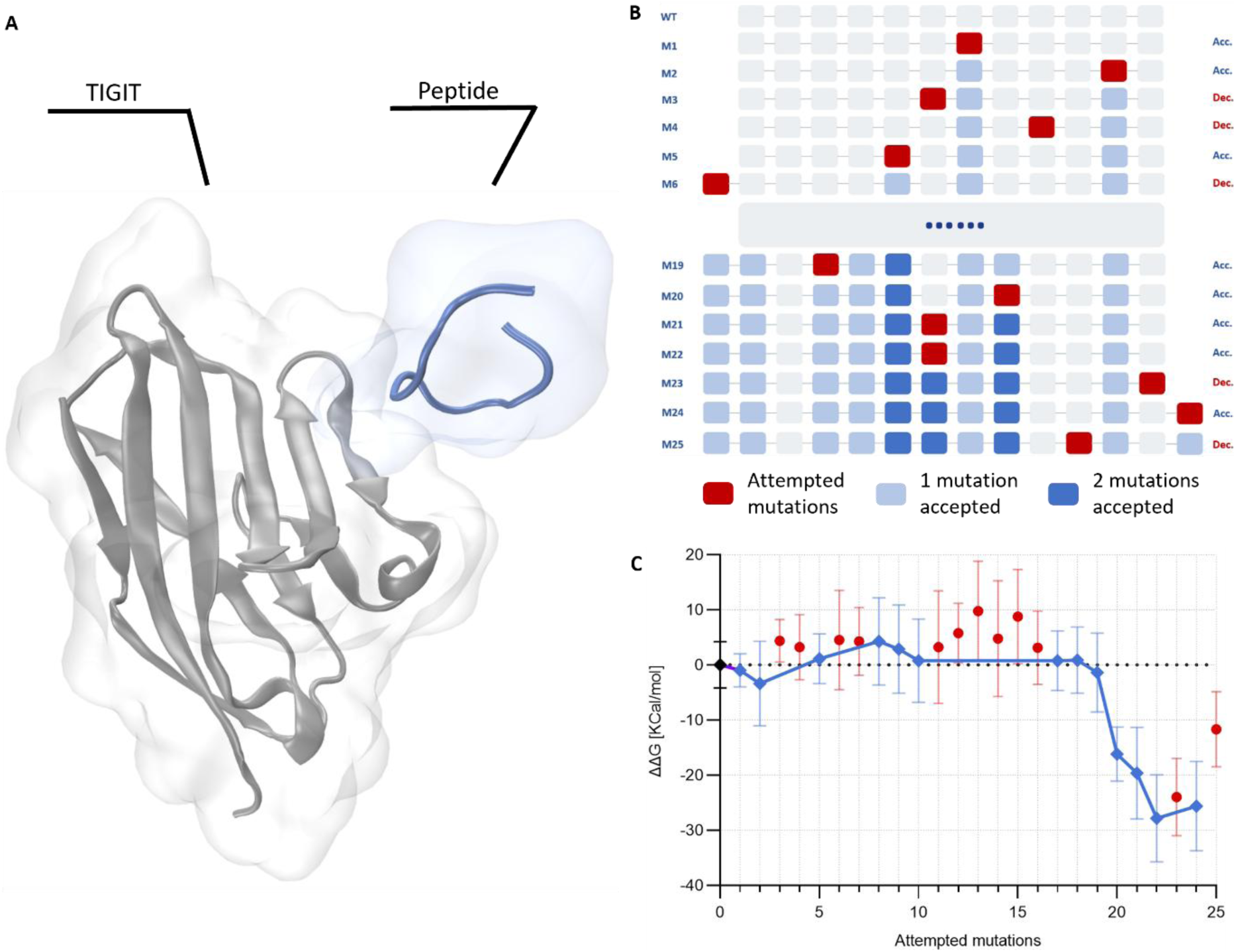
Affinity maturation in peptide mode. The figure shows an example of a Monte Carlo run for the affinity maturation of a peptide against TIGIT. **(A)** The model used for simulations was selected among several models of candidate peptides docked to TIGIT. The initial configuration of the peptide is shown in blue, and the extracellular domain of the TIGIT protein is shown in gray. **(B)** Schematic representations of the positions of attempted and accepted mutations. Each square represents an amino acid belonging to the paratope. The red squares represent sites where mutations are attempted, and darker blue squares represent accepted mutations. Some sites can be mutated more than once. Acc. and Dec. indicate whether the mutation is accepted or declined at that specific step. **(C)** Graph of the binding energy difference along the simulation. Blue points represent the binding energy difference between the wild type and the mutations that have been accepted. The bars indicate the standard error of the computational results.

### Computational efficiency

The examples shown in Figure 4 and Figure 5 are typical runs and demonstrate that the Monte Carlo sampling of the sequence space is very effective in finding sequences that improve the predicted binding affinity in a relatively short number of steps. For the antibody example, we performed 35 different Monte Carlo runs with 30 attempted moves each and obtained 22 sequences with a predicted binding free energy improvement of >15 Kcal/mol in the process (**Figure 6 and Supplementary Figure 1**). While some of these sequences could produce false positives, due to the inherent variance of MMPBSA calculations, our previous experience and preliminary experimental evidence suggest that ANUBI can reliably produce improved binders. A comprehensive validation of the methodology, together with experimental evidence, will be provided in future work.

**Figure 6.**
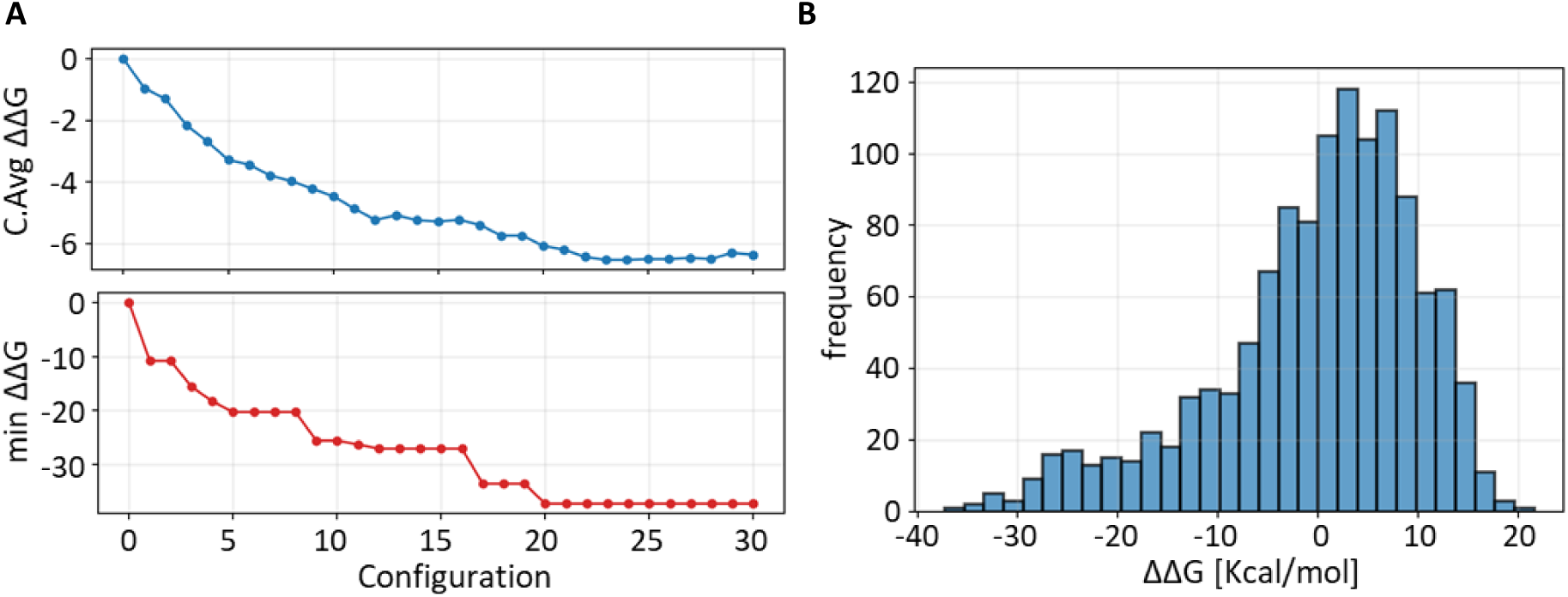
Statistical overview of ANUBI runs for the Cx43 antibody complex. **(A)** Improvement of binding energy along 35 different Monte Carlo runs. The top panel shows the average binding energy improvement as a function of the Monte Carlo step. The bottom panel shows the best binders across all 35 runs as a function of the Monte Carlo step. Individual traces are shown in **Supplementary Figure 1**. **(B)** Histogram of the ΔΔG values for every attempted Monte Carlo step across all 35 runs.

The limiting step in the pipeline is the evaluation of the binding free energy with the MMPBSA method. As shown in Figure 2, the strategy chosen consists of performing ten simulations of 5 ns each and using this ensemble to extract configurations to calculate the binding energy. On a single modern GPU, depending on the size of the system, this process requires 4-10 hours; therefore, we can evaluate several new binders per day. Overall, a typical Monte Carlo run, as shown in Figures 4 and 5, required 1-2 weeks on a single GPU node. Furthermore, the process is highly parallelizable, and many Monte Carlo Markov Chains can be run simultaneously, depending on the computational availability. This timeframe is comparable with analogous experimental maturation of an initial hit, but potentially at a much-reduced cost, depending on the availability of computational resources.

## DISCUSSION

The use of computational methods in biotechnology and drug design has increased constantly in recent years. Beyond the recent expansion of machine learning approaches ^37^, traditional methods based on realistic physical models, such as MD simulations, are benefiting from increased computational power availability. Their ability to accurately sample the configuration space of macromolecules in thermal equilibrium makes them irreplaceable tools for understanding biological properties and in drug development.

In our previous work, we demonstrated that once-expensive binding affinity calculations can now be performed within reasonable timeframes, permitting sensible sampling of the sequence space ^17–19^. This methodology can be used to screen and refine existing antibodies to identify strong binders to a given target ^38^. Similar approaches, paired with emerging *de novo* protein design methods ^16^, can produce virtual pipelines capable of substituting the initial fundamental steps in drug design ^39^.

However, correctly setting up and running such simulations still requires considerable expertise. To lower this entry barrier, we developed ANUBI, the software package presented in this work. ANUBI is designed to be accessible to researchers across different backgrounds, requiring minimal input to initiate automated sequence optimization protocols aimed at improving the binding of proteins or peptides to a desired target. The modular architecture of ANUBI allows users to customize individual components according to their specific needs or computational resources, while maintaining the overall automated workflow.

The critical ingredients for the success of such a pipeline are an accurate calculation of the binding energy and a fast and sensitive sampling of the protein or peptide sequence space.

Crucially, calculation of binding energy must be carried out by averaging the results over several sampled configurations. Proteins and peptides are flexible molecules, with a high number of degrees of freedoms, and the binding energy changes substantially even with minor changes of the side chains positions in space. Configuration sampling in ANUBI is achieved through MD simulations. While relatively expensive from the computational point of view, such sampling provide more reliable estimate of the binding energy and open the way to use ANUBI also for treat flexible targets such as the peptide example discussed here and, possibly in future, intrinsically disordered proteins ^40^ or RNA aptamers ^41^.

Binding energy calculations are then carried out using the MMPBSA method, which is widely adopted as it requires less extensive calculations than alchemical methods ^38,42,43^. Faster methods, based on empirical scoring functions ^44,45^, have also been considered, but they showed poor correlation with experimental data, at least in some specific cases (*R*^2^ = 0.07 for PRODIGY, *R*^2^ = 0.01 for RosettaFF, to be compared with *R*^2^ = 0.57 for the MMPBSA calculation on the same dataset ^18,19^). The major limitation of MMPBSA is the large intrinsic variance of the computed results, as they strongly depend on the configuration of the amino acids that are involved in the binding. To mitigate this effect, we repeat the calculation on ten independent MD trajectories and take the average as the best estimate of the binding energy. We note that MMPBSA does not explicitly calculate entropic contributions to binding. By averaging over equilibrium MD trajectories, we partially account for conformational entropy effects in the bound state through thermal sampling of accessible configurations. We recently demonstrated that, despite this limitation, the MMPBSA method is effective in distinguishing good binders from very good binders, and can therefore be utilized efficiently for screening ^19^.

The sequence space is sampled by a Monte Carlo Metropolis algorithm, which attempts to optimize the binding affinity of the protein or peptide to its target. This method has proven to be consistently successful in identifying predicted improved sequences in a small number of attempts (typically, tens of iterations), on different molecular systems, as shown in **Figures 4, 5 and 6**. Such a sampling can be obtained in 1-3 weeks on a single modern GPU card, a time comparable to that required by affinity maturation experiments, but with a much reduced monetary cost. Moreover, the process is completely parallelizable, depending on the availability of computing power. Many Monte Carlo cycles can be run at the same time, effectively increasing the number of sampled sequences as they diverge very fast from each other, even if they start from the same sequence.

Success in the computational pipeline does not automatically translate into experimental success, and users should be aware of several important limitations. As noted above, MMPBSA’s inability to capture entropy changes upon binding limits the accuracy of absolute affinity predictions, though relative ranking of similar variants remains reliable. Additionally, the homology modeling approach may fail to predict significant conformational changes triggered by certain mutations, particularly those affecting backbone geometry or inducing allosteric effects beyond the immediate binding interface. Furthermore, ANUBI focuses solely on binding affinity and does not evaluate developability criteria essential for therapeutic candidates—including aggregation propensity, solubility, thermal stability, expression yield, or immunogenicity. Mutations may also alter the overall folded state or biophysical properties such as hydrophobicity and toxicity, risks common to all computational design methods that require experimental validation. Future enhancements to the ANUBI platform could address some of these limitations. FEP calculations could be integrated as an additional refinement step for top-ranked candidates, providing more accurate binding affinity estimates that include entropic contributions. For peptide-based therapeutics, computational filters for aggregation propensity ^46^ and toxicity ^47^ could be incorporated. These additions would further reduce the experimental burden by filtering out candidates with poor developability properties before synthesis and testing.

Nevertheless, in our previously published work, two out of three predicted sequences were indeed better binders than the starting one ^18^. Preliminary unpublished data show similar results on different systems. These results demonstrate ANUBI’s practical utility as a refinement tool, effectively enriching for improved variants and significantly reducing the experimental search space compared to traditional affinity maturation approaches. This positions ANUBI as a powerful screening method that can substantially decrease the number of required experimental tests for obtaining a lead molecule in pharmaceutical applications.

## CONCLUSIONS

In this work, we have developed ANUBI, an automated computational pipeline for the affinity maturation of proteins and peptides. This tool is designed to serve as a powerful follow-up step, refining promising candidates generated from either de novo design platforms or initial experimental screenings. By streamlining the simulation workflow, ANUBI aims to democratize the use of advanced molecular modeling, making these powerful predictive methodologies accessible to researchers without requiring extensive computational expertise.

Future work will focus on expanding ANUBI’s capabilities by providing additional options for both energy calculation and sequence sampling modules. More rigorous methods for binding energy calculations, such as FEP, will be offered as an alternative or complementary approach to MMPBSA for cases requiring higher accuracy, while generative AI methods will be integrated as additional options to enhance the efficiency of sequence space exploration. This modular expansion will allow users to tailor the pipeline to their specific requirements while maintaining ANUBI’s ease of use.

## METHODS

### Dependencies

ANUBI relies on the following key components and libraries for full functionality:

### Molecular Dynamics and Free Energy Calculation Software

#### GROMACS (GPU version)

Required for performing molecular dynamics simulations.

#### gmx_MMPBSA

Utilized for MMPBSA binding free energy calculations compatible with GROMACS trajectories.

#### Modeller

Employed for structural mutation modelling.

### Additional Python Libraries

Several Python packages are required for data handling and configuration parsing, including Pandas, NumPy, PyYAML, Biopython. These can be installed via Conda.

The software depends on the tools and libraries and is designed to be fully compatible with them. This compatibility ensures smooth integration into existing workflows. Such seamless interoperability enhances efficiency and user experience during computational tasks.

### Models’ preparation

ANUBI requires an input model of target – peptide or target – antibody interaction, which needs to be ready for MD simulations. Here we report the detailed description of how we built the models described in Figure 4 and Figure 6, as examples, while analogous information regarding the model generation for the TIGIT (Figure 5) will be given in future works.

The starting model for the experimental validation of the antibody against Cx43 was derived from the previously published model of the AbEC1.1-Cx26HC complex ^32,33^. The model was adapted by mutating two critical amino acids in the extracellular (EC) loops to their corresponding Cx43 residues (Q56L in EC1 and H176N in EC2), using MODELLER ^29^. To improve the speed of the simulations, only the EC loop regions (residues 36-78 and 151-195) of the Connexin were retained. To preserve the correct quaternary structure of the system, harmonic positional restraints (300 KJ/mol ·Å^2^) were applied to the heavy atoms of the transmembrane (TM) region residues (36-39, 75-78, 151-154, and 192-195). Such restraints keep the atoms’ position fixed in space, preventing the unfolding of the TM region even in absence of the membrane.

The system was solvated with TIP3P water and neutralized with K⁺ and Cl⁻ ions at 0.15 M to mimic physiological ionic conditions. After solvation, the total number of atoms was approximately 1.3 x 10⁵ atoms. Energy minimization was followed by pre-equilibration in constant volume (NVT) and constant pressure (NPT) ensembles. Specifically, a 250 ps NVT equilibration was carried out with a 1 fs time step, followed by a 1.0 ns NPT equilibration with a 2 fs time step, both under heavy-atom restraints. Production molecular dynamics (MD) simulations were then performed for 50 ns. The starting configuration for ANUBI was extracted from the trajectory after stabilization of the RMSD.

### Automatic modeling during the Monte Carlo cycle

Single amino acids substitutions and N-term or C-term modifications are done with MODELLER ^29^. Since we always start from well-equilibrated models, only the amino acid selected for the modification is modified at each iteration.

This is done by transferring coordinates from the original structure and building missing atoms using internal coordinates derived from bond lengths, angles, and dihedrals. Stereochemical and dihedral restraints are then applied to maintain proper bond geometry and backbone conformation. Finally, local optimization is obtained by combining conjugate gradient minimization with restrained molecular dynamics, in which atomic movements per step are limited to 0.39 Å and the simulation advanced with a time step of 4 fs, allowing the mutated residue and its surrounding environment to relax. During the process, non-bonded interactions are calculated dynamically using the Lennard-Jones potential, considering neighboring atoms within a distance of 4 Å.

The final optimized modified structure is written to a PDB file and used in the follow up analysis.

### MD Simulations, energy calculation and estimation of errors

Periodic boundary conditions are applied to the system generated by MODELLER, and solvation is performed automatically to prepare the system for the following MD simulations, which are carried out in four stages: (i) energy minimization, (ii) 1 ns NVT equilibration with 2 fs time steps and positional restraints on heavy atoms, (iii) 100 ps NPT equilibration with 2 fs time steps, and (iv) and a set of ten 5-ns production MD with 2 fs time steps. During the production MD, the trajectory is saved every 20 ps. The last 150 frames of every production MD are used to compute the binding energy using gmx_MMPBSA ^30^ in a standard linear Poisson-Boltzmann calculation for protein-protein interaction.

Binding free energies are calculated as averages over the 10 independent trajectories (**Supplementary Figure 2**), with reported errors representing standard deviations across replicas. This choice reflects the observation that between-trajectory variance, arising from different sampled configurations of binding interface side chains, dominates over within-trajectory statistical uncertainty. Each trajectory is obtained from the same initial configuration, but with different initial velocities. The length of the different replicas (5 ns) is comparable with the autocorrelation times for side chain rearrangement (2-3 ns ^48,49^) but much longer than the potential energy autocorrelation time (of the order of 1 ps ^50^). This ensures replicas are long enough for reliable energy evaluation while remaining within local conformational basins. This strategy has been proved to be cost effective compared to longer calculations ^19^.

MD simulations in the examples reported above were performed using the GROMACS ^51^ 2023 package, with the Amber99sb-ildn force field ^52^, following protocols previously established ^53^. Long-range electrostatic interactions were treated using the fast smooth particle-mesh Ewald method ^54^ with a standard cut-off of 1.0 nm. Temperature and pressure were maintained at 310 K and 1 atm using the v-rescale and c-rescale coupling algorithms, respectively.

### Randomization and reproducibility

ANUBI relies on stochastic processes at multiple stages of the workflow, and employs the following seeding policy:

**Monte Carlo proposal step:** Mutations are selected randomly from the user-defined list of allowed residues at each position.

**Metropolis acceptance test:** A configuration is accepted if a uniformly distributed random number (0 to 1) is below the computed acceptance probability.

**MODELLER:** Stochastic variation is introduced during side-chain placement and local structural refinement. The random seed is generated via Python’s built-in random module.

**MD initialization:** Initial velocities for each of the 10 replica trajectories are assigned randomly by GROMACS at the start of each simulation.

By default, ANUBI assigns a fresh random seed to each Monte Carlo cycle and Metropolis evaluation. GROMACS automatically randomizes the seed for initial velocity generation unless explicitly specified by the user. This ensures that independent runs explore different regions of sequence and conformational space, even when starting from identical initial configurations.

## CODE AVAILABILITY AND LICENSING

ANUBI is licensed under the GNU Affero General Public License v3.0 (AGPL-3.0) with the Commons Clause Condition v1.0.

The Python code for ANUBI is available at the following github link: https://github.com/ZontaLab/ANUBI

## Founding sources

This work was partially supported by Xi’an Jiaotong-Liverpool University: Research Development Fund RDF-23-01-026 to FZ, Post Graduate Research Project Scholarship FOSA2406006 to WW and FZ; and by Zhejiang University: Fundamental Research Funds for the Central University (226-2024-00123) to DB, and by the National Key R&D Program of China (2024YFA1306400, 2021YFA1201200, 2024YFA1307500), the National Center of Technology Innovation for Biopharmaceuticals (NCTIB2022HS02010), Shanghai Artificial Intelligence Lab (P22KN00272), the Starry Night Science Fund of Zhejiang University Shanghai Institute for Advanced Study (SN-ZJU-SIAS-003) to RZ.

Simulations were carried on at the Xi’an Jiaotong-Liverpool University HPC and the Institute of Quantitative Biology of Zhejiang University HPC.

## Supplementary material file

**Supplementary Figure 1.**
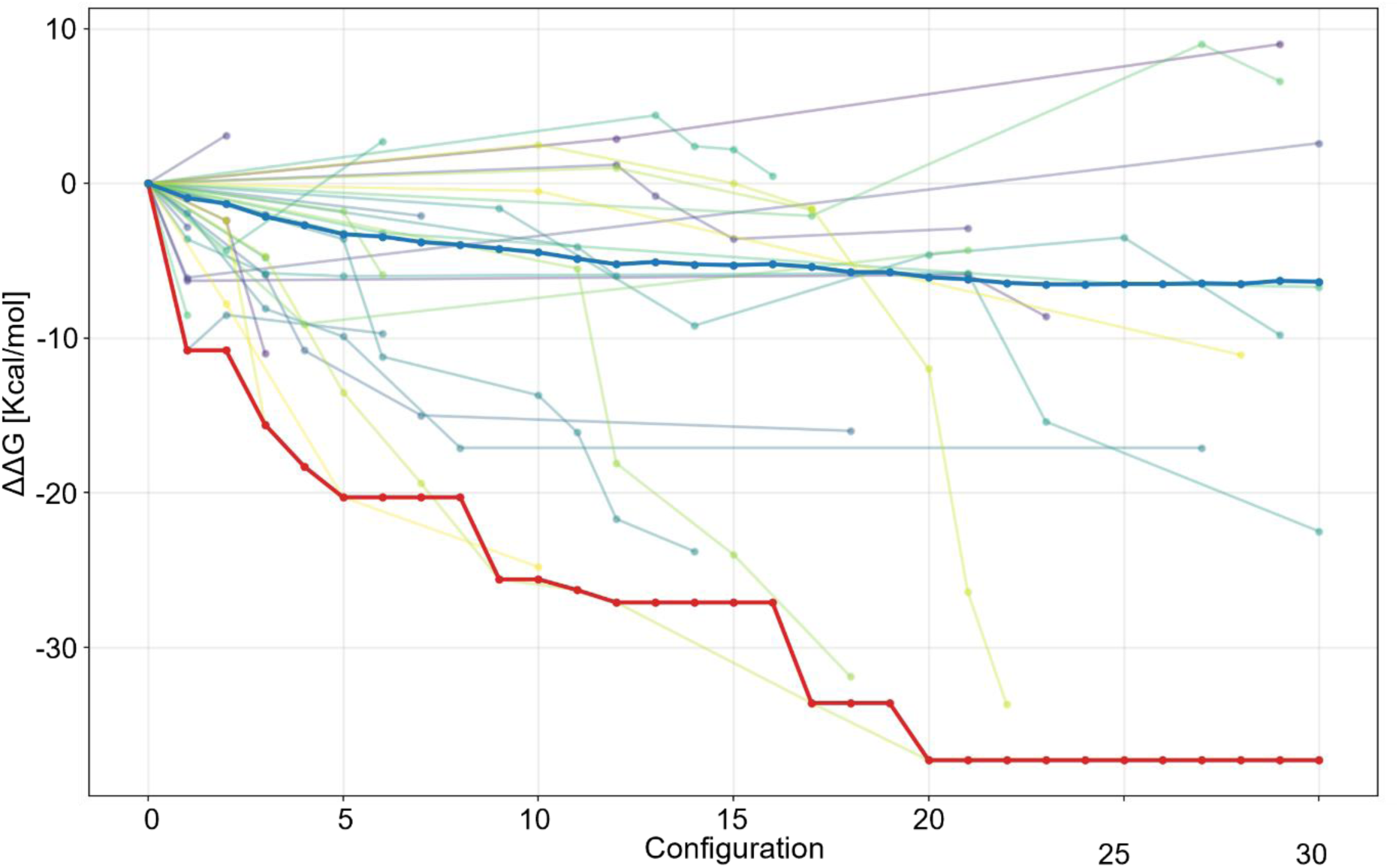
Individual traces for different Monte Carlo runs used for Figure 6. The figure reports 35 Monte Carlo runs for optimization of the antibody binding to the Cx43 hemichannel. For clarity, only accepted modifications are shown. Average and best binders are shown in thick blue line and thick red line, respectively, and are the same as in Panel A of Figure 6. The binding energy can increase in some cases, according to the Metropolis algorithm, but in general, average and best binding affinities rapidly decrease at each iteration.

**Supplementary Figure 2.**
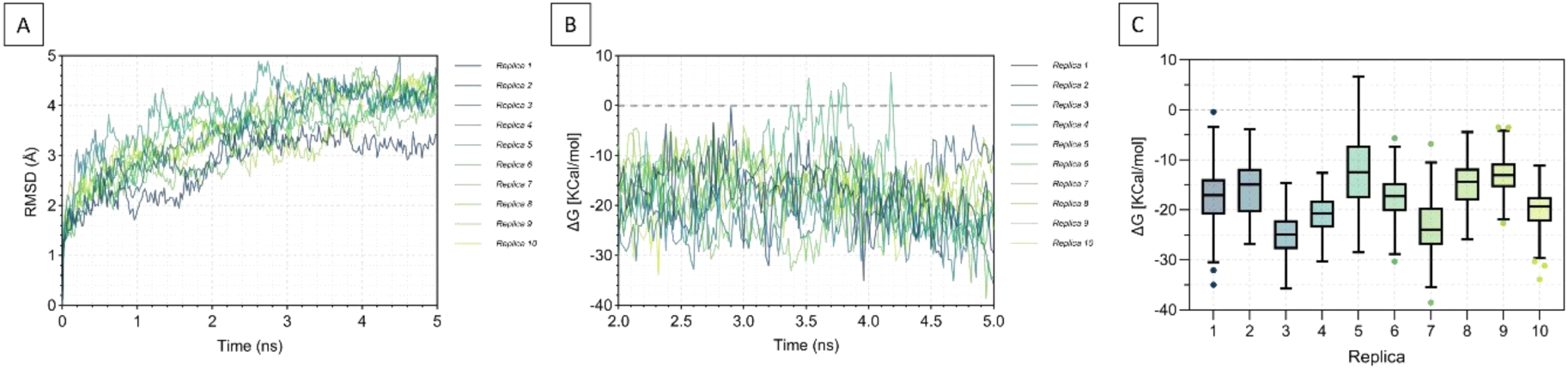
Representative example of MD trajectories in binding energy calculation. (A) RMSD trace for the 10 replicas. Backbone RMSD (nm) versus time for each replica (same alignment as in the main text). Traces are shown over the full 0–5 ns trajectory; the 2–5 ns window was used for quantitative analyses in panels B–C. (B) Binding energy evaluated for the 10 replicas. Binding energy is calculated using the MMPBSA method for each replica over the 2–5 ns interval. (C) Distribution of binding energy for each replica. Data are shown using box-and-whisker plots of the energy for each replica. Boxes show median and IQR; whiskers denote Tukey; outliers are plotted individually. Energies are calculated from configurations extracted from the trajectories of panels A and B (2-5 ns interval only).

## REFERENCES

(1) Bassani, D.; Moro, S. Past, Present, and Future Perspectives on Computer-Aided Drug Design Methodologies. Molecules 2023, 28 (9), 3906. 10.3390/molecules28093906.

(2) Wang, H. Prediction of Protein–Ligand Binding Affinity via Deep Learning Models. Brief Bioinform 2024, 25 (2). 10.1093/bib/bbae081.

(3) Mobley, D. L.; Gilson, M. K. Predicting Binding Free Energies: Frontiers and Benchmarks. Annual Review of Biophysics 2017, 46 (1), 531–558. 10.1146/annurev-biophys-070816-033654.

(4) Eberhardt, J.; Santos-Martins, D.; Tillack, A. F.; Forli, S. AutoDock Vina 1.2.0: New Docking Methods, Expanded Force Field, and Python Bindings. Journal of Chemical Information and Modeling 2021. 10.1021/acs.jcim.1c00203.

(5) Adolf-Bryfogle, J.; Kalyuzhniy, O.; Kubitz, M.; Weitzner, B. D.; Hu, X.; Adachi, Y.; Schief, W. R.; Jr, R. L. D. RosettaAntibodyDesign (RAbD): A General Framework for Computational Antibody Design. PLOS Computational Biology 2018, 14 (4), e1006112. 10.1371/journal.pcbi.1006112.

(6) Friesner, R. A.; Banks, J. L.; Murphy, R. B.; Halgren, T. A.; Klicic, J. J.; Mainz, D. T.; Repasky, M. P.; Knoll, E. H.; Shelley, M.; Perry, J. K.; Shaw, D. E.; Francis, P.; Shenkin, P. S. Glide: A New Approach for Rapid, Accurate Docking and Scoring. 1. Method and Assessment of Docking Accuracy. J. Med. Chem. 2004, 47 (7), 1739–1749. 10.1021/jm0306430.

(7) Passaro, S.; Corso, G.; Wohlwend, J.; Reveiz, M.; Thaler, S.; Somnath, V. R.; Getz, N.; Portnoi, T.; Roy, J.; Stark, H.; Kwabi-Addo, D.; Beaini, D.; Jaakkola, T.; Barzilay, R. Boltz-2: Towards Accurate and Efficient Binding Affinity Prediction. bioRxiv June 18, 2025, p 2025.06.14.659707. 10.1101/2025.06.14.659707.

(8) Bitencourt-Ferreira, G.; de Azevedo, W. F. Machine Learning to Predict Binding Affinity. In Docking Screens for Drug Discovery; de Azevedo Jr., W. F., Ed.; Springer: New York, NY, 2019; pp 251–273. 10.1007/978-1-4939-9752-7_16.

(9) Yang, Y.; Yao, K.; Repasky, M. P.; Leswing, K.; Abel, R.; Shoichet, B. K.; Jerome, S. V. Efficient Exploration of Chemical Space with Docking and Deep Learning. J. Chem. Theory Comput. 2021, 17 (11), 7106–7119. 10.1021/acs.jctc.1c00810.

(10) Abramson, J.; Adler, J.; Dunger, J.; Evans, R.; Green, T.; Pritzel, A.; Ronneberger, O.; Willmore, L.; Ballard, A. J.; Bambrick, J.; Bodenstein, S. W.; Evans, D. A.; Hung, C.-C.; O’Neill, M.; Reiman, D.; Tunyasuvunakool, K.; Wu, Z.; Žemgulytė, A.; Arvaniti, E.; Beattie, C.; Bertolli, O.; Bridgland, A.; Cherepanov, A.; Congreve, M.; Cowen-Rivers, A. I.; Cowie, A.; Figurnov, M.; Fuchs, F. B.; Gladman, H.; Jain, R.; Khan, Y. A.; Low, C. M. R.; Perlin, K.; Potapenko, A.; Savy, P.; Singh, S.; Stecula, A.; Thillaisundaram, A.; Tong, C.; Yakneen, S.; Zhong, E. D.; Zielinski, M.; Žídek, A.; Bapst, V.; Kohli, P.; Jaderberg, M.; Hassabis, D.; Jumper, J. M. Accurate Structure Prediction of Biomolecular Interactions with AlphaFold 3. Nature 2024, 630 (8016), 493–500. 10.1038/s41586-024-07487-w.

(11) Zonta, F.; Pantano, S. From Sequence to Mechanobiology? Promises and Challenges for AlphaFold 3. Mechanobiology in Medicine 2024, 2 (3), 100083. 10.1016/j.mbm.2024.100083.

(12) Honorato, R. V.; Trellet, M. E.; Jiménez-García, B.; Schaarschmidt, J. J.; Giulini, M.; Reys, V.; Koukos, P. I.; Rodrigues, J. P. G. L. M.; Karaca, E.; van Zundert, G. C. P.; Roel-Touris, J.; van Noort, C. W.; Jandová, Z.; Melquiond, A. S. J.; Bonvin, A. M. J. J. The HADDOCK2.4 Web Server for Integrative Modeling of Biomolecular Complexes. Nat Protoc 2024, 19 (11), 3219–3241. 10.1038/s41596-024-01011-0.

(13) Grosdidier, A.; Zoete, V.; Michielin, O. SwissDock, a Protein-Small Molecule Docking Web Service Based on EADock DSS. Nucleic Acids Res 2011, 39 (suppl_2), W270–W277. 10.1093/nar/gkr366.

(14) Kozakov, D.; Hall, D. R.; Xia, B.; Porter, K. A.; Padhorny, D.; Yueh, C.; Beglov, D.; Vajda, S. The ClusPro Web Server for Protein-Protein Docking. Nat Protoc 2017, 12 (2), 255–278. 10.1038/nprot.2016.169.

(15) Lisanza, S. L.; Gershon, J. M.; Tipps, S. W. K.; Sims, J. N.; Arnoldt, L.; Hendel, S. J.; Simma, M. K.; Liu, G.; Yase, M.; Wu, H.; Tharp, C. D.; Li, X.; Kang, A.; Brackenbrough, E.; Bera, A. K.; Gerben, S.; Wittmann, B. J.; McShan, A. C.; Baker, D. Multistate and Functional Protein Design Using RoseTTAFold Sequence Space Diffusion. Nat Biotechnol 2025, 43 (8), 1288–1298. 10.1038/s41587-024-02395-w.

(16) Pacesa, M.; Nickel, L.; Schellhaas, C.; Schmidt, J.; Pyatova, E.; Kissling, L.; Barendse, P.; Choudhury, J.; Kapoor, S.; Alcaraz-Serna, A.; Cho, Y.; Ghamary, K. H.; Vinué, L.; Yachnin, B. J.; Wollacott, A. M.; Buckley, S.; Westphal, A. H.; Lindhoud, S.; Georgeon, S.; Goverde, C. A.; Hatzopoulos, G. N.; Gönczy, P.; Muller, Y. D.; Schwank, G.; Swarts, D. C.; Vecchio, A. J.; Schneider, B. L.; Ovchinnikov, S.; Correia, B. E. One-Shot Design of Functional Protein Binders with BindCraft. Nature 2025, 1–10. 10.1038/s41586-025-09429-6.

(17) Buratto, D.; Saxena, A.; Ji, Q.; Yang, G.; Pantano, S.; Zonta, F. Rapid Assessment of Binding Affinity of SARS-COV-2 Spike Protein to the Human Angiotensin-Converting Enzyme 2 Receptor and to Neutralizing Biomolecules Based on Computer Simulations. Frontiers in Immunology 2021, 12.

(18) Buratto, D.; Wan, Y.; Shi, X.; Yang, G.; Zonta, F. In Silico Maturation of a Nanomolar Antibody against the Human CXCR2. Biomolecules 2022, 12 (9), 1285. 10.3390/biom12091285.

(19) Autiero, I.; Buratto, D.; Guo, F.; Wang, W.; Biswal, M. R.; Chan, K. C.; Zhou, R.; Zonta, F. Assessing Computational Strategies for the Evaluation of Antibody Binding Affinities. Journal of Chemical Theory and Computation 2025. 10.1021/acs.jctc.5c01231.

(20) Zheng, B.; Wang, X.; Guo, M.; Tzeng, C.-M. Therapeutic Peptides: Recent Advances in Discovery, Synthesis, and Clinical Translation. Int J Mol Sci 2025, 26 (11), 5131. 10.3390/ijms26115131.

(21) Genheden, S.; Ryde, U. The MM/PBSA and MM/GBSA Methods to Estimate Ligand-Binding Affinities. Expert Opinion on Drug Discovery 2015, 10 (5), 449–461. 10.1517/17460441.2015.1032936.

(22) Xiao, W.; Jiang, W.; Chen, Z.; Huang, Y.; Mao, J.; Zheng, W.; Hu, Y.; Shi, J. Advance in Peptide-Based Drug Development: Delivery Platforms, Therapeutics and Vaccines. Sig Transduct Target Ther 2025, 10 (1), 74. 10.1038/s41392-024-02107-5.

(23) Jenkins, G. W.; Safonova, Y.; Smider, V. V. Germline-Encoded Positional Cysteine Polymorphisms Enhance Diversity in Antibody Ultralong CDR H3 Regions. J Immunol 2022, 209 (11), 2141–2148. 10.4049/jimmunol.2200455.

(24) Hayes, R. L.; Brooks, C. L. A Strategy for Proline and Glycine Mutations to Proteins with Alchemical Free Energy Calculations. J Comput Chem 2021, 42 (15), 1088–1094. 10.1002/jcc.26525.

(25) Aguilar-Gurrieri, C.; Barajas, A.; Rovirosa, C.; Ortiz, R.; Urrea, V.; de la Iglesia, N.; Clotet, B.; Blanco, J.; Carrillo, J. Alanine-Based Spacers Promote an Efficient Antigen Processing and Presentation in Neoantigen Polypeptide Vaccines. Cancer Immunol Immunother 2023, 72 (7), 2113–2125. 10.1007/s00262-023-03409-3.

(26) Wang, G.; Li, X.; Wang, Z. APD3: The Antimicrobial Peptide Database as a Tool for Research and Education. Nucleic Acids Res 2016, 44 (D1), D1087–1093. 10.1093/nar/gkv1278.

(27) Marqus, S.; Pirogova, E.; Piva, T. J. Evaluation of the Use of Therapeutic Peptides for Cancer Treatment. Journal of Biomedical Science 2017, 24 (1), 21. 10.1186/s12929-017-0328-x.

(28) O’Brien, C.; Flower, D. R.; Feighery, C. Peptide Length Significantly Influences in Vitro Affinity for MHC Class II Molecules. Immunome Res 2008, 4, 6. 10.1186/1745-7580-4-6.

(29) Webb, B.; Sali, A. Comparative Protein Structure Modeling Using MODELLER. Curr Protoc Bioinformatics 2016, 54, 5.6.1-5.6.37. 10.1002/cpbi.3.

(30) Valdés-Tresanco, M. S.; Valdés-Tresanco, M. E.; Valiente, P. A.; Moreno, E. gmx_MMPBSA: A New Tool to Perform End-State Free Energy Calculations with GROMACS. J. Chem. Theory Comput. 2021, 17 (10), 6281–6291. 10.1021/acs.jctc.1c00645.

(31) Zhou, R.; Das, P.; Royyuru, A. K. Single Mutation Induced H3N2 Hemagglutinin Antibody Neutralization: A Free Energy Perturbation Study. J. Phys. Chem. B 2008, 112 (49), 15813–15820. 10.1021/jp805529z.

(32) Xu, L.; Carrer, A.; Zonta, F.; Qu, Z.; Ma, P.; Li, S.; Ceriani, F.; Buratto, D.; Crispino, G.; Zorzi, V.; Ziraldo, G.; Bruno, F.; Nardin, C.; Peres, C.; Mazzarda, F.; Salvatore, A. M.; Raspa, M.; Scavizzi, F.; Chu, Y.; Xie, S.; Yang, X.; Liao, J.; Liu, X.; Wang, W.; Wang, S.; Yang, G.; Lerner, R. A.; Mammano, F. Design and Characterization of a Human Monoclonal Antibody That Modulates Mutant Connexin 26 Hemichannels Implicated in Deafness and Skin Disorders. Frontiers in Molecular Neuroscience 2017, 10.

(33) Ziraldo, G.; Buratto, D.; Kuang, Y.; Xu, L.; Carrer, A.; Nardin, C.; Chiani, F.; Salvatore, A. M.; Paludetti, G.; Lerner, R. A.; Yang, G.; Zonta, F.; Mammano, F. A Human-Derived Monoclonal Antibody Targeting Extracellular Connexin Domain Selectively Modulates Hemichannel Function. Frontiers in Physiology 2019, 10.

(34) Buratto, D.; Donati, V.; Zonta, F.; Mammano, F. Harnessing the Therapeutic Potential of Antibodies Targeting Connexin Hemichannels. Biochimica et Biophysica Acta (BBA) - Molecular Basis of Disease 2021, 1867 (4), 166047. 10.1016/j.bbadis.2020.166047.

(35) Jeong, B.-S.; Nam, H.; Lee, J.; Park, H.-Y.; Cho, K. J.; Sheen, J. H.; Song, E.; Oh, M.; Lee, S.; Choi, H.; Yang, J.-E.; Kim, M.; Oh, B.-H. Structural and Functional Characterization of a Monoclonal Antibody Blocking TIGIT. MAbs 2022, 14 (1), 2013750. 10.1080/19420862.2021.2013750.

(36) Stengel, K. F.; Harden-Bowles, K.; Yu, X.; Rouge, L.; Yin, J.; Comps-Agrar, L.; Wiesmann, C.; Bazan, J. F.; Eaton, D. L.; Grogan, J. L. Structure of TIGIT Immunoreceptor Bound to Poliovirus Receptor Reveals a Cell–Cell Adhesion and Signaling Mechanism That Requires Cis-Trans Receptor Clustering. Proc Natl Acad Sci U S A 2012, 109 (14), 5399–5404. 10.1073/pnas.1120606109.

(37) Noé, F.; Tkatchenko, A.; Müller, K.-R.; Clementi, C. Machine Learning for Molecular Simulation. Annual Review of Physical Chemistry 2020, 71 (1), 361–390. 10.1146/annurev-physchem-042018-052331.

(38) Roux, B.; Chipot, C. Editorial Guidelines for Computational Studies of Ligand Binding Using MM/PBSA and MM/GBSA Approximations Wisely. J. Phys. Chem. B 2024, 128 (49), 12027–12029. 10.1021/acs.jpcb.4c06614.

(39) Watson, J. L.; Juergens, D.; Bennett, N. R.; Trippe, B. L.; Yim, J.; Eisenach, H. E.; Ahern, W.; Borst, A. J.; Ragotte, R. J.; Milles, L. F.; Wicky, B. I. M.; Hanikel, N.; Pellock, S. J.; Courbet, A.; Sheffler, W.; Wang, J.; Venkatesh, P.; Sappington, I.; Torres, S. V.; Lauko, A.; De Bortoli, V.; Mathieu, E.; Ovchinnikov, S.; Barzilay, R.; Jaakkola, T. S.; DiMaio, F.; Baek, M.; Baker, D. De Novo Design of Protein Structure and Function with RFdiffusion. Nature 2023, 620 (7976), 1089–1100. 10.1038/s41586-023-06415-8.

(40) Muhammedkutty, F. N. K.; MacAinsh, M.; Zhou, H.-X. Atomistic Molecular Dynamics Simulations of Intrinsically Disordered Proteins. Current Opinion in Structural Biology 2025, 92, 103029. 10.1016/j.sbi.2025.103029.

(41) Gram, H.; Marconi, L. A.; Barbas, C. F.; Collet, T. A.; Lerner, R. A.; Kang, A. S. In Vitro Selection and Affinity Maturation of Antibodies from a Naive Combinatorial Immunoglobulin Library. Proceedings of the National Academy of Sciences 1992, 89 (8), 3576–3580. 10.1073/pnas.89.8.3576.

(42) Aldeghi, M.; Heifetz, A.; J. Bodkin, M.; Knapp, S.; C. Biggin, P. Accurate Calculation of the Absolute Free Energy of Binding for Drug Molecules. Chemical Science 2016, 7 (1), 207–218. 10.1039/C5SC02678D.

(43) Shivakumar, D.; Williams, J.; Wu, Y.; Damm, W.; Shelley, J.; Sherman, W. Prediction of Absolute Solvation Free Energies Using Molecular Dynamics Free Energy Perturbation and the OPLS Force Field. J. Chem. Theory Comput. 2010, 6 (5), 1509–1519. 10.1021/ct900587b.

(44) Xue, L. C.; Rodrigues, J. P.; Kastritis, P. L.; Bonvin, A. M.; Vangone, A. PRODIGY: A Web Server for Predicting the Binding Affinity of Protein–Protein Complexes. Bioinformatics 2016, 32 (23), 3676–3678. 10.1093/bioinformatics/btw514.

(45) Alford, R. F.; Leaver-Fay, A.; Jeliazkov, J. R.; O’Meara, M. J.; DiMaio, F. P.; Park, H.; Shapovalov, M. V.; Renfrew, P. D.; Mulligan, V. K.; Kappel, K.; Labonte, J. W.; Pacella, M. S.; Bonneau, R.; Bradley, P.; Dunbrack, R. L.; Das, R.; Baker, D.; Kuhlman, B.; Kortemme, T.; Gray, J. J. The Rosetta All-Atom Energy Function for Macromolecular Modeling and Design. Journal of Chemical Theory and Computation 2017, 13 (6), 3031–3048. 10.1021/acs.jctc.7b00125.

(46) Barrera, E. E.; Zonta, F.; Pantano, S. Dissecting the Role of Glutamine in Seeding Peptide Aggregation. Computational and Structural Biotechnology Journal 2021, 19, 1595–1602. 10.1016/j.csbj.2021.02.014.

(47) Rathore, A. S.; Choudhury, S.; Arora, A.; Tijare, P.; Raghava, G. P. S. ToxinPred 3.0: An Improved Method for Predicting the Toxicity of Peptides. Comput Biol Med 2024, 179, 108926. 10.1016/j.compbiomed.2024.108926.

(48) Sapienza, P. J.; Lee, A. L. Using NMR to Study Fast Dynamics in Proteins: Methods and Applications. Curr Opin Pharmacol 2010, 10 (6), 723–730. 10.1016/j.coph.2010.09.006.

(49) Lindorff-Larsen, K.; Piana, S.; Palmo, K.; Maragakis, P.; Klepeis, J. L.; Dror, R. O.; Shaw, D. E. Improved Side-Chain Torsion Potentials for the Amber ff99SB Protein Force Field. Proteins: Structure, Function, and Bioinformatics 2010, 78 (8), 1950–1958. 10.1002/prot.22711.

(50) Iwai, R.; Kasahara, K.; Takahashi, T. Influence of Various Parameters in the Replica-Exchange Molecular Dynamics Method: Number of Replicas, Replica-Exchange Frequency, and Thermostat Coupling Time Constant. Biophysics and Physicobiology 2018, 15, 165–172. 10.2142/biophysico.15.0_165.

(51) Pronk, S.; Páll, S.; Schulz, R.; Larsson, P.; Bjelkmar, P.; Apostolov, R.; Shirts, M. R.; Smith, J. C.; Kasson, P. M.; van der Spoel, D.; Hess, B.; Lindahl, E. GROMACS 4.5: A High-Throughput and Highly Parallel Open Source Molecular Simulation Toolkit. Bioinformatics 2013, 29 (7), 845–854. 10.1093/bioinformatics/btt055.

(52) Maier, J. A.; Martinez, C.; Kasavajhala, K.; Wickstrom, L.; Hauser, K. E.; Simmerling, C. ff14SB: Improving the Accuracy of Protein Side Chain and Backbone Parameters from ff99SB. Journal of Chemical Theory and Computation 2015, 11 (8), 3696–3713. 10.1021/acs.jctc.5b00255.

(53) Nielsen, B. S.; Zonta, F.; Farkas, T.; Litman, T.; Nielsen, M. S.; MacAulay, N. Structural Determinants Underlying Permeant Discrimination of the Cx43 Hemichannel. Journal of Biological Chemistry 2019, 294 (45), 16789–16803. 10.1074/jbc.RA119.007732.

(54) Darden, T.; York, D.; Pedersen, L. Particle Mesh Ewald: An N⋅log(N) Method for Ewald Sums in Large Systems. J. Chem. Phys. 1993, 98 (12), 10089–10092. 10.1063/1.464397.

